# Effects of increased N and P availability on biomass allocation and root carbohydrate reserves differ between N-fixing and non-N-fixing savanna tree seedlings

**DOI:** 10.1101/224188

**Authors:** Varan Varma, Arockia M Catherin, Mahesh Sankaran

## Abstract

In mixed tree-grass ecosystems, tree recruitment is limited by demographic bottlenecks to seedling establishment arising from inter- and intra-life form competition, and disturbances such as fire. Enhanced nutrient availability resulting from anthropogenic nitrogen (N) and phosphorus (P) deposition can alter the nature of these bottlenecks by changing seedling growth and biomass allocation patterns, and lead to longer-term shifts in tree community composition if different plant functional groups respond differently to increased nutrient availability. However, the extent to which tree functional types characteristic of savannas differ in their responses to increased N and P availability remains unclear. We quantified differences in above- and belowground biomass, and root carbohydrate contents – parameters known to influence the ability of plants to compete, as well as survive and recover from fires – in seedlings of multiple N-fixing and non-N-fixing tree species characteristic of Indian savanna and dry-forest ecosystems to experimental N and P additions. N-fixers in our study were co-limited by N and P availability, while non-N-fixers were N limited. Although both functional groups increased biomass production following fertilisation, non-N-fixers were more responsive and showed greater relative increases in biomass with fertilisation than N-fixers. N-fixers had greater baseline investment in belowground resources and root carbohydrate stocks, and while fertilisation reduced root:shoot ratios in both functional groups, root carbohydrate content only reduced with fertilisation in non-N-fixers. Our results indicate that, even within a given system, plants belonging to different functional groups can be limited by, and respond differentially to, different nutrients, suggesting that long-term consequences of nutrient deposition are likely to vary across savannas contingent on the relative amounts of N and P being deposited in sites.

## Introduction

The structure and functioning of mixed tree-grass ecosystems such as savannas is governed by both bottom-up (e.g. water and nutrient availability) and top-down drivers (e.g. fire and herbivory) (Frost et al. 1986, Scholes and Archer 1997, Higgins et al. 2000, Sankaran et al. 2004, 2008, February et al. 2007, Ekblom and Gillson 2010). Amongst bottom-up drivers, the importance of water availability in regulating savanna structure and dynamics is well recognised (Sankaran et al. 2005, Lehmann et al. 2014). However, the influence of nutrient availability on the functioning of these ecosystems is less clear (Sankaran et al. 2008, February and Higgins 2010, van der Waal et al. 2011). Given that savannas and dry forest ecosystems are anticipated to be particularly vulnerable to future global change drivers including nutrient deposition (Sala et al. 2000), understanding the impacts of enhanced nutrient availability on vegetation dynamics of mixed tree-grass systems is important, both to assess their future trajectories and to develop appropriate management strategies.

Atmospheric nutrient deposition, a major global change driver, has dramatically increased the quantities of plant available nitrogen (N) and phosphorus (P) cycling through ecosystems across the globe (Vitousek 1994, Vitousek et al. 1997, Bennett et al. 2001, Galloway et al. 2004, 2008, Phoenix et al. 2006). This increased availability of N and P has the potential to impact both the structure and composition of mixed tree-grass communities. For example, in African savannas, tree basal area has been shown to be negatively correlated with soil N across broad scales (Sankaran et al. 2008), suggesting that enhanced N availability can potentially alter vegetation structure by shifting communities towards more grassy states. Nutrient deposition can also affect the composition of tree communities if increased nutrient availability has differing effects on the dominant tree functional types that characterise these ecosystems, namely, N-fixers and non-N-fixers.

N-fixing and non-N-fixing species differ inherently in their nutritional requirements, leaf chemistry and physiology (Vitousek et al. 2002, 2013, Pearson and Vitousek 2002, Powers and Tiffin 2010), and thus may be expected to respond differently to increases in N and P availability (Khurana and Singh 2004, Cramer et al. 2007, 2010, Barbosa et al. 2014). By virtue of their association with N-fixing bacteria in root nodules, N-fixing plants are characterised by an N and P demanding lifestyle compared to non-N-fixers (Vitousek et al. 2002, 2013, Pearson and Vitousek 2002). Nodulation, though an advantage in N limited soils, incurs a substantial energetic cost on the plant (Gutschick 1981, Vitousek and Howarth 1991, Sprent 1999, Vitousek et al. 2002, 2010), and studies have previously shown that N-fixers reduce investment in root nodules with increasing N availability (Sanginga et al. 1988, Barron et al. 2010). Reduced investment in nodulation under conditions of increased nutrient availability can free up additional resources that can be diverted towards growth in N-fixers. However, whether or not such effects translate to greater biomass increases in N-fixers relative to non-N-fixers that have a lower N and P demand, is unclear.

Besides altering growth and allocation patterns, increased nutrient availability can also influence how tree seedlings respond to fire, a major determinant of vegetation structure in mixed tree-grass ecosystems. The ability of both tree seedlings and juveniles to survive recurring fires by resprouting is key to their persistence and eventual recruitment into the canopy as reproductively mature adults (Bond and van Wilgen 1996, Higgins et al. 2000, Sankaran et al. 2004, 2005, Bond 2008, Hanan et al. 2008, Schutz et al. 2009). This capacity to resprout, and hence, persist within the fire trap is contingent on seedlings being able to allocate sufficient resources below ground and invest in root carbohydrate reserves, which are remobilised to support the cost of post-fire resprouting (Ryle et al. 1981, Pate et al. 1990, Bell and Ojeda 1999, Hoffmann et al. 2000, Bell 2001, Verdaguer and Ojeda 2002, Bond et al. 2003, Lamont and Wiens 2003, Vesk and Westoby 2004, Knox and Clarke 2005, Schwilk and Ackerly 2005, Hermans et al. 2006, Clarke and Knox 2009, Wigley et al. 2009, Clarke et al. 2013, Wang et al. 2015). Allocation of resources below ground has been shown to be influenced by nutrient availability (Tilman 1988, Knox and Clarke 2005, Hermans et al. 2006, Wang et al. 2015), and plants in general tend to reduce below ground investment, i.e. lower root-shoot ratios (henceforth, R:S ratios), and decrease root carbohydrate reserves with increasing N and P availability (Ryle et al. 1981, Hermans et al. 2006, Clarke and Knox 2009, Wang et al. 2015). Differences between species and functional groups in allocation to below ground resources and root carbohydrate reserves with increasing nutrient availability can significantly alter post-fire survival and the composition of the regenerating community, but few studies have thus far evaluated how N-fixers and non-N-fixers differ in their allocation patterns with N and P addition.

In this study, we quantified the effects of N and P fertilisation on biomass accumulation, biomass partitioning and root storage carbohydrate content in seedlings of multiple tree species characteristic of savanna and tropical dry forests in peninsular India. We chose these responses as they play an important role in determining the competitive ability of tree seedlings, as well as their ability to survive fires, a key disturbance agent in this ecosystem. We examined how responses differed between N-fixers and non-N-fixers, and additionally, quantified changes in nodulation in N-fixers when fertilised. We expected: a) enhanced nutrient availability to lead to increased biomass accumulation in both functional groups, but the magnitude of increase to be greater amongst non-N-fixers, b) N-fixers to have greater baseline R:S ratios, as well as root carbohydrate content compared to non-N-fixers, and c) nutrient addition to reduce R:S ratios and root carbohydrate content in both functional groups, with greater reductions in non-N-fixers.

## Materials & Methods

The experiment was conducted at a field site in the village of Hosur, located in Mysore district of the southern Indian state of Karnataka. A total of 13 commonly occurring savanna and tropical dry forest tree species were selected based on published sources (Puyravaud et al. 1994, Sagar and Singh 2004, Kumar and Shahabuddin 2005, Kodandapani et al. 2008) and included six N-fixers (*Acacia catechu* (L.f.) Willd., *Acacia ferruginea* DC., *Acacia leucophloea* (Roxb.) Willd., *Albizia amara* (Roxb.) B.Biovin., *Albizia lebbeck* (L.) Benth. an*d Dalbergia latifolia* (Roxb.)) and seven non-N-fixers (*Lagerstroemia indica* (L.), *Lagerstroemia speciosa* (L.) Pers., *Phyllanthus emblica* (L.), *Sapindus emarginatus* Vahl., *Terminalia arjuna* (Roxb. ex DC.), *Terminalia bellirica* (Gaertn.) Roxb. and *Zizyphus jujuba* Mill.).

Three week old seedlings were procured from the Foundation for the Revitalisation of Local Health and Tradition (FRLHT), Bangalore, transported to the field site and allowed to acclimatise to local conditions for one week, and then transplanted individually in to 20 L nursery polybags containing a 1:1 mixture of sand and local soil (July 2013). Average total C and N content of the sand-soil mix was 3.94 g.Kg^−1^ and 0.51 g. Kg^−1^, respectively (Leco TrueSpec CN analyser), and average total P content was 0.13 g.Kg^−1^ (Thermo iCAP 6300 ICP – OES dual view spectrophotometer). Plants were placed randomly in a uniform grid with a minimum spacing of 40 cm between individual stems. Each individual was assigned to one of four nutrient treatments – Control (Con; no nutrient addition), N+ (5g N), P+ (0.5g P) and NP+ (5g N and 0.5g P). N and P were added to polybags as solutions of urea and single superphosphate (SSP), respectively, in three separate applications two, four and six weeks after transplant. Seedlings were watered regularly to prevent water stress. Final sample sizes for each species-treatment combination ranged from five to 14 individuals, with a total of 624 individuals in the entire experiment.

All individuals were harvested six months after transplant (January, 2014). Shoots and roots were separated, and for the N-fixers, root nodules were collected. Samples were transported to the National Centre for Biological Sciences, Bangalore, where they were oven dried for 5 days at 60°C before being weighed. At the time of weighing root samples, a section of the primary root from a subset of individuals, ranging from four to six individuals per species-treatment combination, was extracted to estimate percent storage carbohydrates, i.e. non-structural carbohydrates (NSC), using the phenol-sulphuric acid assay (Buysse and Merckx 1993, Wigley et al. 2009). Ground root samples were digested in 3% HCl, followed by the addition of 27% phenol and concentrated sulphuric acid to the supernatant of the acid digest. The intensity of the resulting colour reaction was measured using a spectrophotometer at 490 nm and sample carbohydrate content was estimated against a standard curve of glucose.

Responses of seedling total-, above- and belowground biomass, R:S ratios and root carbohydrate content to fertilisation were analysed using linear mixed effects models, implemented using the *lme4* package (Bates et al. 2014) in R (R Core Team 2014). Predictor variables included nutrient treatment, plant functional group and the interaction between the two, with species identity included as a random factor as the focus of the analyses were to find generalisations at the functional group level, while accounting for intrinsic differences between species in the measured parameters. Significance tests of the fixed effects were carried out using Satterthwaite's approximation for degrees of freedom implemented within the *lmerTest* package (Kuznetova et al. 2014). The nodule mass data for N-fixers included a large number of zeros for the nutrient addition treatments resulting in a skewed distribution which did not match criteria to be considered a zero inflated distribution. Hence, for this analysis, we used species-treatment means of nodule mass as the response variable within a linear mixed effects model, where nutrient treatment was the only predictor, and species identity included as a random factor.

## Results

### Biomass accumulation and allocation

N-fixers and non-N-fixers differed in their response to nutrient addition (significant nutrient treatment x functional group interaction for total biomass: *F* = 5.6984, *df* = 3, *P* < 0.001; shoot biomass: *F* = 6.1188, *df* = 3, *P* < 0.001; root biomass: *F* = 4.7438, *df* = 3, *P* = 0.003). N-fixers in this system appeared to be co-limited by N and P, showing significant increases in total (55%; *P* < 0.001; Fig 1a), shoot (64%; *P* = 0.001; Fig 1c) and root biomass (51%; *P* < 0.001; Fig 1e) only when supplied with both N and P. Non-N-fixers, on the other hand, appeared to be N limited, with increases in total (63%; *P* < 0.001; Fig 1b), shoot (91%; *P* < 0.001; Fig 1d) and root biomass (46%; *P* < 0.001; Fig 1f) observed for the N addition treatment, which were virtually identical to the increases observed in the NP+ treatment (59%, 83% and 45%, for total, shoot and root biomass, respectively). P addition had no effect on biomass responses.

**Fig. 1.**
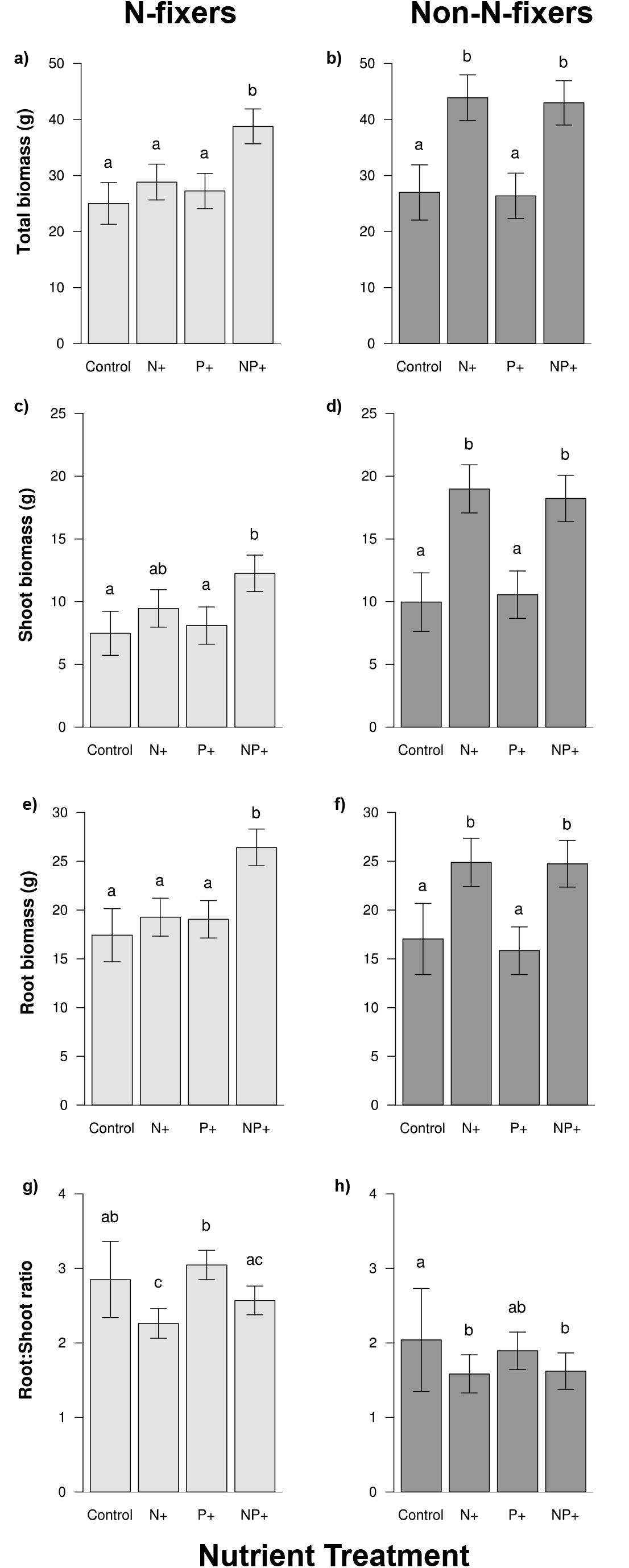
Response of N-fixing and non-N-fixing species to N and P fertilisation with respect to total biomass (a,b), aboveground biomass (c,d), belowground biomass accumulation (e,f) and root-shoot (R:S) ratios (g,h). Letters indicate significant differences within functional groups. Error bars represent 1 SE.

Differences in the responses of N-fixers and non-N-fixers to nutrient addition were strongest for shoot biomass. N-fixers increased their above-ground biomass by 64% from approximately 7.47 g in the control treatment to 12.25 g in the NP+ treatment (Fig 1c), while biomass in non-N-fixers increased by 83% from 9.96 g to 18.22 g (Fig 1d). Similarly, total plant biomass increased by 55% from 25 g in the control treatment to 38.77 g in the NP+ treatment for N-fixers (Fig 1a), while for the same treatment combinations, total plant biomass increased by 59% from 26.98 g to 42.97 g in non-N-fixers (Fig 1b). Increases in root biomass with nutrient addition were of similar magnitude for both functional groups (Fig 1e,f).

On average, N-fixers tended to invest relatively more in below-ground tissue (R:S = 2.8, SE = 0.51) compared to non-N-fixers (R:S = 2.04, SE = 0.69), although this difference was not significant (P = 0.27; Fig 1g,h). For both N-fixers and non-N-fixers, R:S ratios declined in response to nutrient addition (nutrient treatment x functional group interaction: NS). However, within functional groups, N-fixer (Fig 1g) R:S ratios declined by 21% with N addition (P = 0.003) and by 10% in the NP+ treatment (NS), while non-N-fixers (Fig 1h) demonstrated significant declines in both the N+ (-22%; *P* = 0.004) and NP+ treatments (-21%; *P* = 0.007).

### Root carbohydrate storage

N-fixers had significantly greater concentrations of root carbohydrates compared to non-N-fixers (*P* = 0.003, Fig 2). None of the nutrient treatments had any effect on root carbohydrate concentrations of N-fixing species. In contrast, root carbohydrate concentrations in non-N-fixers declined significantly relative to controls following N addition, both when applied alone (N+) and in combination with P (i.e. NP+). P addition did not affect root carbohydrate concentrations (Fig 2b). For non-N-fixers, root storage carbohydrate contents declined from 12.6% in the controls to 10.6% in the N+ (P = 0.01) and NP+ (P = 0.01) treatments; a reduction of 16%.

**Fig 2.**
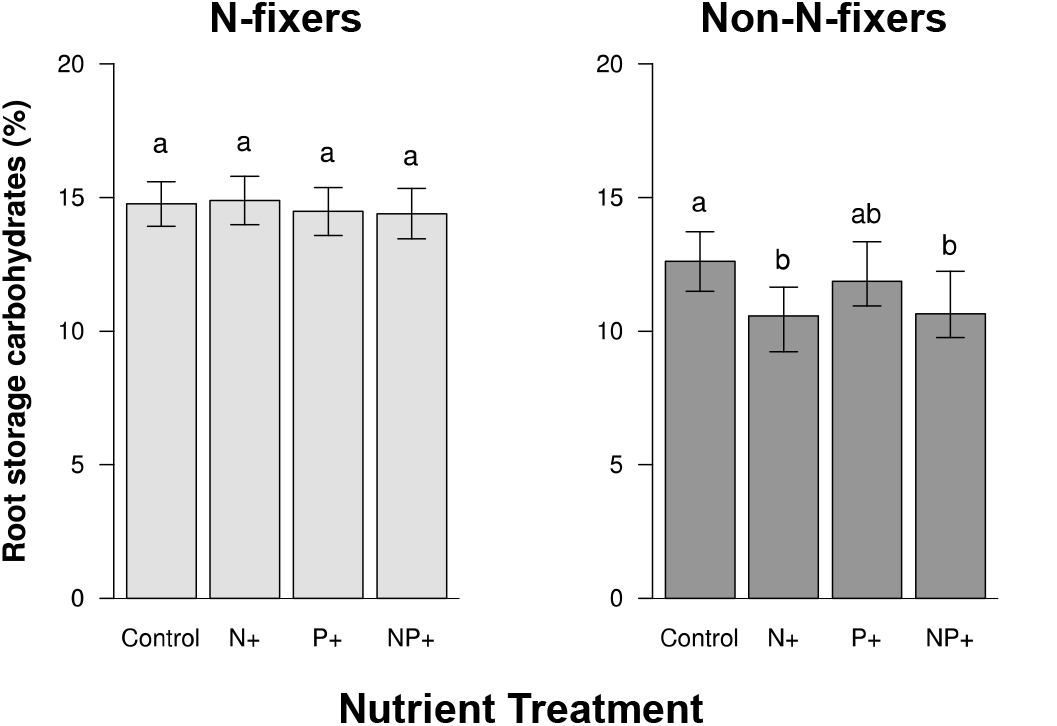
Modification in percent root storage carbohydrates in response to N and P addition for N-fixing and non-N-fixing species. Letters indicate significant differences within functional groups. Error bars represent 1

### Nodulation in N-fixers

All N-fixing species nodulated in the control treatment. Fertilisation had a very strong negative effect on total nodule mass (species means) in N-fixers (*F* = 4.6622, *df* = 3, *P* = 0.02; Fig 3). Combined N and P addition resulted in the largest declines (-88%; *P* = 0.004) in nodulation, followed by N addition (-67%; *P* = 0.02). P addition resulted in marginal, but non-significant reductions nodule weight.

**Fig 3.**
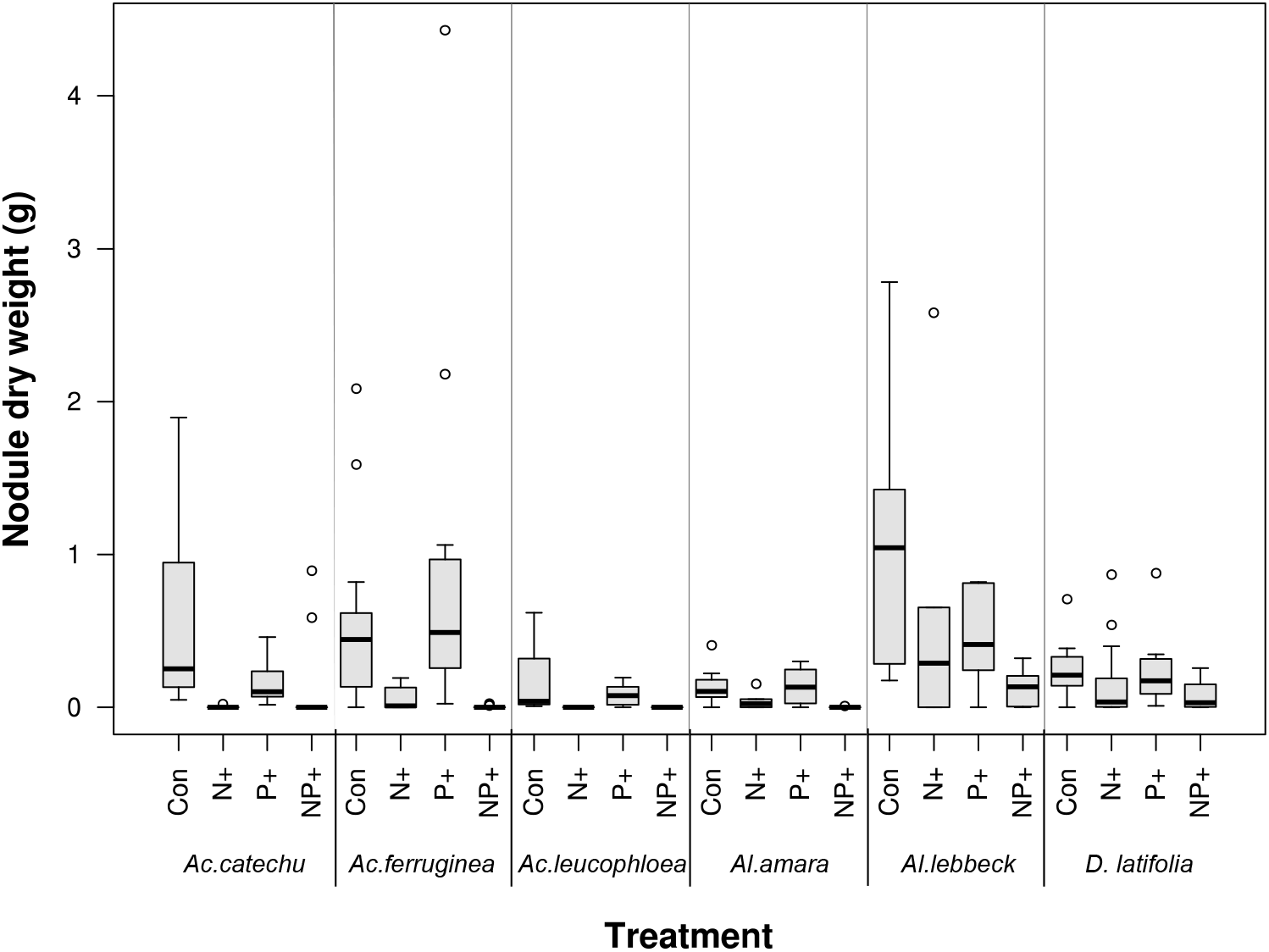
Changes in nodule dry weight in N-fixing species in response to N and P addition.

## Discussion

Our results indicate that differences in the nature of nutrient limitation between N-fixing and non-N-fixing tropical dry forest tree seedlings can lead to contrasting responses of the two functional groups to atmospheric nutrient deposition. Although seedlings of both functional groups increased biomass when fertilised, N-fixers increased biomass only when simultaneously supplied with both N and P, while non-N-fixers responded only to N addition. Further, the magnitude of biomass increase with fertilisation was greater for non-N-fixers when compared to N-fixers. Baseline allocation in root biomass and root storage carbohydrates was greater in N-fixers, and though both functional groups tended to reduce relative investment in root biomass with nutrient addition, only non-N-fixers showed concurrent reductions in root storage carbohydrate content.

N-fixers and non-N-fixers in our study differed in the nature of their nutrient limitation, with growth of non-N-fixing species limited by N availability, while N-fixers were co-limited by N and P in accordance with their N and P demanding lifestyle (Vitousek et al. 2002, 2013, Pearson and Vitousek 2002). Previous studies that have investigated the nature of nutrient limitation of savanna vegetation have largely tended to focus on the herbaceous component of savannas (Ludwig et al. 2001, Barger et al. 2002, Craine et al. 2008, Ries and Shugart 2008, Cech et al. 2008, O’Halloran et al. 2010, Copeland et al. 2012, Bustamante et al. 2012). Studies that have considered savanna trees have typically evaluated woody vegetation responses to the addition of a single nutrient (Kraaij and Ward 2006, Wang et al. 2012), or the combined addition of N and P (Khurana and Singh 2004, van Der Waal et al. 2009, Vadigi and Ward 2013, Barbosa et al. 2014), thereby precluding identification of the specific nutrient(s) limiting growth (but see – Wang et al. 2012, Holdo 2013). Results from these earlier studies suggest contrasting patterns of nutrient limitation of herbaceous vegetation across the diverse savannas of the world, ranging from nutrients not being limiting (O’Halloran et al. 2010), to grass growth being N-limited (Cech et al. 2008, Wang et al. 2012), P-limited (Ludwig et al. 2001), or co-limited by N and P (Craine et al. 2008, Cech et al. 2008). Woody plant responses, where evaluated, have also been similarly varied with studies reporting no effects of N addition on growth of a non-N-fixing species (*Colophospermum mopane;* van Der Waal et al. 2009), P-limitations to growth of an N-fixing species (*Acacia erioloba;* Wang et al. 2012), and possible N and P co-limitation of growth of a non-N-fixing species (*Combretum hereroense;* Holdo 2013). There is also evidence to suggest that different life-forms within a given savanna can be limited by different nutrients. For example, in a Namibian savanna, grass growth was reported to be N-limited while trees displayed P-limitation (Wang et al. 2012). While there have been surprisingly few studies that have investigated whether N-fixing and non-N-fixing savanna trees differ in the nature of their nutrient limitation, results from this study and inferences based on differing N:P ratios of savannah grasses, N-fixing and non-N-fixing trees (Ratnam et al. 2008, Pellegrini 2016) suggest that differential nutrient limitation of different plant functional groups in savannas may potentially be a common phenomenon. Ultimately, our results suggest that the effects of nutrient deposition on vegetation dynamics of mixed tree-grass ecosystems is likely to be contingent on the relative rates of N and P being deposited in sites, with responses varying both spatially and between plant functional types within a site.

Non-N-fixers in our study were more responsive to the alleviation of nutrient limitation and showed greater relative increases in biomass with fertilisation compared to N-fixers. Although N-fixers significantly reduced root nodule mass with increasing N availability (N and NP+ treatments), as expected given the energetic costs of N fixation (Gutschick 1981, Sterner and Elser 2002, Vitousek et al. 2002), and potentially diverted some of these resources to growth (Davidson and Robson 1986, Boakye et al. 2015), the magnitude of biomass increase was not as pronounced as that observed in the case of non-N-fixers. These results are consistent with broader scale patterns reported in a meta-analysis by Xia and Wan (2008), where biomass increases in non-N-fixers were twice as large as those of N-fixers across a range of plant life forms and ecosystem types. Differences in responses to fertilisation between functional groups were particularly pronounced for shoot biomass, with non-N-fixers nearly doubling aboveground biomass (83% increase) with combined N and P addition, as opposed to a 64% increase in N-fixers. Barbosa et al. (2014) also report similar results, where NPK fertilisation resulted in increases in seedling stem length only amongst non-N-fixing South African savanna tree species. Greater relative investment in aboveground growth can be an effective strategy for juveniles to avoid light competition and quickly escape the zone of grass fuelled fires above which fire-induced tree mortality is low (i.e. the fire trap), when fires are infrequent. However, it potentially comes at the cost of being able to survive, resprout and persist within the fire trap when fires are frequent. Although both plant functional types in our study increased total investment in both above- and belowground tissues following fertilisation, non-N-fixers reduced relative investment in belowground tissues (R:S ratios) to a greater extent than N-fixers. Further, non-N-fixers reduced root carbohydrate stocks following fertilisation, while N-fixers adopted a more ‘conservative’ strategy and continued to maintain their root carbohydrate reserves. Greater investment in belowground tissues (R:S ratios), and root carbohydrate stocks in particular, have been linked to faster rates of post-fire recovery (Hoffmann et al. 2000, Bell 2001, Bond et al. 2003, Lamont and Wiens 2003, Vesk and Westoby 2004, Clarke and Knox 2009, Wigley et al. 2009, Clarke et al. 2013) suggesting that N-fixing species are likely to be less prone to fire-mediated mortality following fertilisation. The extent to which enhanced nutrient availability drives long-term shifts in woody plant communities is, therefore, likely to vary spatially contingent on local fire regimes.

Differences observed between N-fixers and non-N-fixers in our study in terms of investment in root carbohydrate reserves following fertilisation can potentially be attributed to differences in the type of reserves that the two functional groups invest in, i.e. true and accumulated reserves (Chapin et al. 1990). True reserves represent baseline plant investment in carbohydrate reserves, whose formation trades-off with allocation of resources to plant growth, maintenance and reproduction, and is not influenced by changes in resource availability (Knox and Clarke 2005, Clarke and Knox 2009). Accumulated reserves, on the other hand, are formed in addition to true reserves when the acquisition of non-limiting resources exceeds demands for growth (Chapin et al. 1990), i.e. when plants are faced with nutrient limitation for growth, but when adequate supplies of other resources are available for photosynthesis to continue. When supplied with limiting nutrients, carbohydrates previously contributing towards the formation of accumulated reserves are utilised to enhance growth. The lack of changes in root carbohydrate content with fertilisation suggests that root carbohydrate reserves in N-fixers may be exclusively made up of true reserves. In contrast, the reductions in percent root storage carbohydrates in non-N-fixers with N addition implies the formation of accumulated reserves in addition to true reserves, which may be associated with the larger increases in biomass production observed in non-N-fixers when fertilised.

Here, we investigated the responses of N-fixing and non-N-fixing savanna and dry-forest woody seedlings to atmospheric nutrient deposition – a pervasive, global change driver, focussing on patterns of seedling growth and biomass allocation when grown alone. While we recognize that ultimate responses are likely to be influenced by factors such as grass competition, we nevertheless believe that our results provide a basis for understanding savanna responses to nutrient deposition. Our results suggest that nutrient deposition has the potential to induce longer-term compositional shifts in savanna tree communities by differentially affecting the growth, post-fire survival and resprouting ability of N-fixing and non-N-fixing species. Further, responses are likely to differ between sites contingent on multiple factors including underlying edaphic characteristics, fire regimes and the relative amounts of N and P being deposited in sites.

## Acknowledgements

We would like to thank Kavita Isvaran, Suhel Quader and Jayashree Ratnam for their input during planning and execution of the experiment. We also thank Dr. Ganesh Babu and Umesh VJ at the Foundation for the Revitalisation of Local Health and Tradition (FRLHT), Bangalore, for providing the tree seedlings, and Manjunatha HC, Meenakshi HJ and Chandregowda J for the use of their land for the experiment. Our gratitude to our field assistants Mahesh HK, Bomrai HK and Mahadev HK, and also to Anil PA, Harinandanan PV, Nandita Natraj, Yadugiri VT, Atul Joshi and Prashath Gowda for their assistance during data collection and lab work. Funding for this work was provided by the National Centre for Biological Sciences (NCBS), Bangalore. This manuscript was greatly improved by comments from Anand M Osuri, Fiona Savory, Sumanta Baghchi and Edmund C February.

## References

Barbosa, E. R. M. et al. 2014. Tree species from different functional groups respond differently to environmental changes during establishment. – Oecologia 174: 1345–1357.

Barger, N. N. et al. 2002. Nutrient limitation to primary productivity in a secondary savanna in Venezuela. – Biotropica 34: 493–501.

Barron, A. R. et al. 2010. Facultative nitrogen fixation by canopy legumes in a lowland tropical forest. – Oecologia 165: 511–520.

Bates, D. et al. 2014. lme4: Linear mixed-effects models using Eigen and S4. – J. Stat. Softw.: ArXiv e-print http://arxiv.org/abs/1406.5823.

Bell, D. T. 2001. Ecological response syndromes in the flora of southwestern Western Australia: Fire resprouters versus reseeders. – Bot. Rev. 67: 417–440.

Bell, T. L. and Ojeda, F. 1999. Underground starch storage in *Erica* species of the Cape Floristic Region – differences between seeders and resprouters. – New Phytol. 144: 143–152.

Bennett, E. M. et al. 2001. Human impact on erodable phosphorus and eutrophication: a global perspective increasing accumulation of phosphorus in soil threatens rivers, lakes, and coastal oceans with eutrophication. – BioScience 51: 227–234.

Boakye, E. Y. et al. 2015. Growth and nodulation response of six indigenous trees and two shrubby legumes to phosphorus and nitrogen fertilizers in two soils of Ghana. – J. Trop. Agric. 53: 21–34.

Bond, W. J. 2008. What limits trees in C_4_ grasslands and savannas? – Annu. Rev. Ecol. Evol. Syst. 39: 641–659.

Bond, W. J. and van Wilgen, B. W. 1996. Fire and plants. – Chapman and Hall.

Bond, W. J. et al. 2003. What controls South African vegetation — climate or fire? – South Afr. J. Bot. 69: 79–91.

Bustamante, M. M. C. et al. 2012. Effects of nutrient additions on plant biomass and diversity of the herbaceous-subshrub layer of a Brazilian savanna (Cerrado). – Plant Ecol. 213: 795–808.

Buysse, J. and Merckx, R. 1993. An improved colorimetric method to quantify sugar content of plant tissue. – J. Exp. Bot. 44: 1627–1629.

Cech, P. G. et al. 2008. Effects of herbivory, fire and N_2_-fixation on nutrient limitation in a humid African savanna. – Ecosystems 11: 991–1004.

Chapin, F. S. et al. 1990. The ecology and economics of storage in plants. – Annu. Rev. Ecol. Syst. 21: 423–447.

Clarke, P. J. and Knox, K. J. E. 2009. Trade-offs in resource allocation that favour resprouting affect the competitive ability of woody seedlings in grassy communities. – J. Ecol. 97: 1374–1382.

Clarke, P. J. et al. 2013. Resprouting as a key functional trait: how buds, protection and resources drive persistence after fire. – New Phytol. 197: 19–35.

Copeland, S. M. et al. 2012. Short-term effects of elevated precipitation and nitrogen on soil fertility and plant growth in a Neotropical savanna. – Ecosphere 3: 1–20.

Craine, J. M. et al. 2008. Nutrient concentration ratios and co-limitation in South African grasslands. – New Phytol. 179: 829–836.

Cramer, M. D. et al. 2007. Grass competition induces N_2_ fixation in some species of African *Acacia*. – J. Ecol. 95: 1123–1133.

Cramer, M. D. et al. 2010. Growth of N_2_-fixing African savanna *Acacia* species is constrained by below-ground competition with grass. – J. Ecol. 98: 156–167.

Davidson, I. A. and Robson, M. J. 1986. Effect of contrasting patterns of nitrate application on the nitrate uptake, N_2_-fixation, nodulation and growth of white clover. – Ann. Bot. 57: 331–338.

Ekblom, A. and Gillson, L. 2010. Hierarchy and scale: testing the long term role of water, grazing and nitrogen in the savanna landscape of Limpopo National Park (Mozambique). – Landsc. Ecol. 25: 1529–1546.

February, E. C. and Higgins, S. I. 2010. The distribution of tree and grass roots in savannas in relation to soil nitrogen and water. – South Afr. J. Bot. 76: 517–523.

February, E. C. et al. 2007. Tree distribution on a steep environmental gradient in an arid savanna. – J. Biogeogr. 34: 270–278.

Frost, P. G. H. et al. 1986. Response of savannas to stress and disturbance. – Biol. Int. Spec. Issue 10 IUBS Paris

Galloway, J. N. et al. 2004. Nitrogen cycles: past, present, and future. – Biogeochemistry 70: 153–226.

Galloway, J. N. et al. 2008. Transformation of the nitrogen cycle: recent trends, questions, and potential solutions. – Science 320: 889–892.

Gutschick, V. P. 1981. Evolved strategies in nitrogen acquisition by plants. – Am. Nat. 118: 607–637.

Hanan, N. P. et al. 2008. Do fires in savannas consume woody biomass? A comment on approaches to modeling savanna dynamics. – Am. Nat. 171: 851–856.

Hermans, C. et al. 2006. How do plants respond to nutrient shortage by biomass allocation? – Trends Plant Sci. 11: 610–617.

Higgins, S. I. et al. 2000. Fire, resprouting and variability: a recipe for grass–tree coexistence in savanna. – J. Ecol. 88: 213–229.

Hoffmann, W. A. et al. 2000. Elevated CO_2_ enhances resprouting of a tropical savanna tree. – Oecologia 123: 312–317.

Holdo, R. M. 2013. Effects of fire history and N and P fertilization on seedling biomass, Specific Leaf Area, and root:shoot ratios in a South African savannah. – South Afr. J. Bot. 86: 5–8.

Khurana, E. and Singh, J. S. 2004. Impact of elevated nitrogen inputs on seedling growth of five dry tropical tree species as affected by life-history traits. – Can. J. Bot. 82: 158–167.

Knox, K. J. E. and Clarke, P. J. 2005. Nutrient availability induces contrasting allocation and starch formation in resprouting and obligate seeding shrubs. – Funct. Ecol. 19: 690–698.

Kodandapani, N. et al. 2008. A comparative analysis of spatial, temporal, and ecological characteristics of forest fires in seasonally dry tropical ecosystems in the Western Ghats, India. – For. Ecol. Manag. 256: 607–617.

Kraaij, T. and Ward, D. 2006. Effects of rain, nitrogen, fire and grazing on tree recruitment and early survival in bush-encroached savanna, South Africa. – Plant Ecol. 186: 235–246.

Kumar, R. and Shahabuddin, G. 2005. Effects of biomass extraction on vegetation structure, diversity and composition of forests in Sariska Tiger Reserve, India. – Environ. Conserv. 32: 248–259.

Kuznetova, A. et al. 2014. lmerTest: Tests for random and fixed effects for linear mixed effect models (lmer objects of lme4 package). R package version 2.0-11.

Lamont, B. B. and Wiens, D. 2003. Are seed set and speciation rates always low among species that resprout after fire, and why? – Evol. Ecol. 17: 277–292.

Lehmann, C. E. R. et al. 2014. Savanna vegetation-fire-climate relationships differ among continents. – Science 343: 548–552.

Ludwig, F. et al. 2001. Effects of nutrients and shade on tree-grass interactions in an East African savanna. – J. Veg. Sci. 12: 579–588.

O’Halloran, L. R. et al. 2010. Nutrient limitations on aboveground grass production in four savanna types along the Kalahari Transect. – J. Arid Environ. 74: 284–290.

Pate, J. S. et al. 1990. Seedling growth and storage characteristics of seeder and resprouter species of Mediterranean-type ecosystems of S.W. Australia. – Ann. Bot. 65: 585–601.

Pearson, H. L. and Vitousek, P. M. 2002. Soil phosphorus fractions and symbiotic nitrogen fixation across a substrate-age gradient in Hawaii. – Ecosystems 5: 587–596.

Pellegrini, A. F. A. 2016. Nutrient limitation in tropical savannas across multiple scales and mechanisms. – Ecology 97: 313–324.

Phoenix, G. K. et al. 2006. Atmospheric nitrogen deposition in world biodiversity hotspots: the need for a greater global perspective in assessing N deposition impacts. – Glob. Change Biol. 12: 470–476.

Powers, J. S. and Tiffin, P. 2010. Plant functional type classifications in tropical dry forests in Costa Rica: leaf habit versus taxonomic approaches. – Funct. Ecol. 24: 927–936.

Puyravaud, J.-P. et al. 1994. Ecotone structure as an indicator of changing forest-savanna boundaries (Linganamakki region, southern India). – J. Biogeogr. 21: 581–593.

R Core Team 2014. R: A language and environment for statistical computing. R Foundation for Statistical Computing, Vienna, Austria. URL http://www.R-project.org/.

Ratnam, J. et al. 2008. Nutrient resorption patterns of plant functional groups in a tropical savanna: variation and functional significance. – Oecologia 157: 141–151.

Ries, L. P. and Shugart, H. H. 2008. Nutrient limitations on understory grass productivity and carbon assimilation in an African woodland savanna. – J. Arid Environ. 72: 1423–1430.

Ryle, G. J. A. et al. 1981. Distribution of dry weight between root and shoot in white clover dependent on N_2_ fixation or utilizing abundant nitrate nitrogen. – Plant Soil 60: 29–39.

Sagar, R. and Singh, J. S. 2004. Local plant species depletion in a tropical dry deciduous forest of northern India. – Environ. Conserv. 31: 55–62.

Sala, O. E. et al. 2000. Global biodiversity scenarios for the year 2100. – Science 287: 1770–1774.

Sanginga, N. et al. 1988. Nodulation and growth of *Leucaena leucocephala* (Lam.) de Wit as affected by inoculation and N fertilizer. – Plant Soil 112: 129–135.

Sankaran, M. et al. 2004. Tree–grass coexistence in savannas revisited – insights from an examination of assumptions and mechanisms invoked in existing models. – Ecol. Lett. 7: 480–490.

Sankaran, M. et al. 2005. Determinants of woody cover in African savannas. – Nature 438: 846–849.

Sankaran, M. et al. 2008. Woody cover in African savannas: the role of resources, fire and herbivory. – Glob. Ecol. Biogeogr. 17: 236–245.

Scholes, R. J. and Archer, S. R. 1997. Tree-grass interactions in savannas. – Annu. Rev. Ecol. Syst. 28: 517–544.

Schutz, A. E. N. et al. 2009. Juggling carbon: allocation patterns of a dominant tree in a fire-prone savanna. – Oecologia 160: 235–246.

Schwilk, D. W. and Ackerly, D. D. 2005. Is there a cost to resprouting? Seedling growth rate and drought tolerance in sprouting and nonsprouting *Ceanothus* (Rhamnaceae). – Am. J. Bot. 92: 404–410.

Sprent, J. I. 1999. Nitrogen fixation and growth of non-crop legume species in diverse environments. – Perspect. Plant Ecol. Evol. Syst. 2: 149–162.

Sterner, R. W. and Elser, J. J. 2002. Ecological stoichiometry: the biology of elements from molecules to the biosphere. – Princeton University Press.

Tilman, D. 1988. Plant strategies and the dynamics and structure of plant communities. – Princeton University Press.

Vadigi, S. and Ward, D. 2013. Shade, nutrients, and grass competition are important for tree sapling establishment in a humid savanna. – Ecosphere 4: art142.

van Der Waal, C. et al. 2009. Water and nutrients alter herbaceous competitive effects on tree seedlings in a semi-arid savanna. – J. Ecol. 97: 430–439.

van der Waal, C. et al. 2011. Large herbivores may alter vegetation structure of semi-arid savannas through soil nutrient mediation. – Oecologia 165: 1095–1107.

Verdaguer, D. and Ojeda, F. 2002. Root starch storage and allocation patterns in seeder and resprouter seedlings of two Cape Erica (*Ericaceae*) species. – Am. J. Bot. 89: 1189–1196.

Vesk, P. A. and Westoby, M. 2004. Funding the bud bank: a review of the costs of buds. – Oikos 106: 200–208.

Vitousek, P. M. 1994. Beyond global warming: ecology and global change. – Ecology 75: 1861–1876.

Vitousek, P. M. and Howarth, R. W. 1991. Nitrogen limitation on land and in the sea: How can it occur? – Biogeochemistry 13: 87–115.

Vitousek, P. M. et al. 1997. Human alteration of the global nitrogen cycle: sources and consequences. – Ecol. Appl. 7: 737–750.

Vitousek, P. M. et al. 2002. Towards an ecological understanding of biological nitrogen fixation. – Biogeochemistry 57-58: 1–45.

Vitousek, P. M. et al. 2010. Terrestrial phosphorus limitation: mechanisms, implications, and nitrogen–phosphorus interactions. – Ecol. Appl. 20: 5–15.

Vitousek, P. M. et al. 2013. Biological nitrogen fixation: rates, patterns and ecological controls in terrestrial ecosystems. – Philos. Trans. R. Soc. B Biol. Sci. 368: 20130119.

Wang, L. et al. 2012. The interactive nutrient and water effects on vegetation biomass at two African savannah sites with different mean annual precipitation. – Afr. J. Ecol. 50: 446–454.

Wang, Y.-L. et al. 2015. Contrasting responses of root morphology and root-exuded organic acids to low phosphorus availability in three important food crops with divergent root traits. – AoB Plants: plv097.

Wigley, B. J. et al. 2009. Sapling survival in a frequently burnt savanna: mobilisation of carbon reserves in *Acacia karroo*. – Plant Ecol. 203: 1–11.

Xia, J. and Wan, S. 2008. Global response patterns of terrestrial plant species to nitrogen addition. – New Phytol. 179: 428–439.

